# Comprehensive comparison of transcriptomes in SARS-CoV-2 infection: alternative entry routes and innate immune responses

**DOI:** 10.1101/2021.01.07.425716

**Authors:** Yingying Cao, Xintian Xu, Simo Kitanovski, Lina Song, Jun Wang, Pei Hao, Daniel Hoffmann

**Affiliations:** Bioinformatics and Computational Biophysics, Faculty of Biology and Center for Medical Biotechnology, University of Duisburg-Essen, Essen 45141, Germany; Key Laboratory of Molecular Virology and Immunology, Institut Pasteur of Shanghai, Center for Biosafety Mega-Science, Chinese Academy of Sciences, Shanghai 200031, China; Translational Skin Cancer Research, German Consortium for Translational Cancer Research, Essen, Germany; The Joint Program in Infection and Immunity: a. Guangzhou Women and Children’s Medical Center, Guangzhou Medical University, Guangzhou 510623, China; b. Institut Pasteur of Shanghai, Chinese Academy of Sciences, Shanghai 200031, China

## Abstract

The pathogenesis of COVID-19 emerges as complex, with multiple factors leading to injury of different organs. Several studies on underlying cellular processes have produced contradictory claims, e.g. on SARS-CoV-2 cell entry or innate immune responses. However, clarity in these matters is imperative for therapy development. We therefore performed a meta-study with a diverse set of transcriptomes under infections with SARS-CoV-2, SARS-CoV and MERS-CoV, including data from different cells and COVID-19 patients. Using these data, we investigated viral entry routes and innate immune responses. First, our analyses support the existence of cell entry mechanisms for SARS and SARS-CoV-2 other than the ACE2 route with evidence of inefficient infection of cells without expression of ACE2; expression of TMPRSS2/TPMRSS4 is unnecessary for efficient SARS-CoV-2 infection with evidence of efficient infection of A549 cells transduced with a vector expressing human ACE2. Second, we find that innate immune responses in terms of interferons and interferon simulated genes are strong in relevant cells, for example Calu3 cells, but vary markedly with cell type, virus dose, and virus type.

## Introduction

Coronaviruses are non-segmented positive-sense RNA viruses with a genome of around 30 kilobases. The genome has a 5’ cap structure along with a 3’ poly (A) tail, which acts as mRNA for translation of the replicase polyproteins. The replicase gene occupies approximately two thirds of the entire genome and encodes 16 non-structural proteins (nsps). The remaining third of the genome contains open reading frames (orfs) that encode accessory proteins and four structural proteins, including spike (S), envelope (E), membrane (M), and nucleocapsid (N) (*1*).

Over the past 20 years, three epidemics or pandemics of life-threatening diseases have been caused by three closely related coronaviruses – severe acute respiratory syndrome coronavirus (SARS-CoV), which emerged with nearly 10 % mortality (*2, 3*) in 2002-2003 and spread to 26 countries before being contained; Middle East respiratory syndrome coronavirus (MERS-CoV), with mortality around 34 % (*4, 5*) starting in 2012 and since then spreading to 27 countries; SARS-CoV-2, emerging in late 2019 (*6*), which has caused many millions of confirmed cases and > 1 million deaths worldwide (*7*). Infection with SARS-CoV, MERS-CoV or SARS-CoV-2 can cause a severe acute respiratory illness with similar symptoms, including fever, cough, and shortness of breath.

SARS-CoV-2 is a new coronavirus, but its similarity to SARS-CoV (amino acid sequences about 76% identical (*8*)) and MERS-CoV suggests comparisons to these earlier epidemics. Despite the difference in the total number of cases caused by SARS-CoV and SARS-CoV-2 (*3, 7*) due to different transmission rates, the outbreak caused by SARS-CoV-2 resembles the outbreak of SARS: both emerged in winter and were linked to exposure to wild animals sold at markets. Although MERS-CoV has high morbidity and mortality rates, lack of autopsies from MERS-CoV cases has hindered our understanding of MERS-CoV pathogenesis in humans.

Until now there are no specific anti-SARS-CoV-2, anti-SARS-CoV or anti-MERS-CoV therapeutics approved for human use. There are several points of attack for potential anti-SARS-CoV-2/SARS-CoV/MERS-CoV therapies, e.g. intervention on cell entry mechanisms to prevent virus invasion, or acting on the host immune system to kill the infected cells and thus prevent replication of the invading viruses. A better understanding of virus entry mechanisms and the immune responses can therefore guide the development of novel therapeutics.

Virus entry into host cells is the first step of the viral life cycle. It is an essential component of cross-species transmission and an important determinant of virus pathogenesis and infectivity (*9, 10*), and also constitutes an antiviral target for treatment and prevention (*11*). It seems that SARS-CoV and SARS-CoV-2 use similar virus entry mechanisms (*12*). The infection of SARS-CoV or SARS-CoV-2 in target cells was initially identified to occur by cell-surface membrane fusion (*13,14*). Some later studies have shown that SARS-CoV can infect cells through receptor mediated endocytosis (*15, 16*) as well. Both mechanisms require the S protein of SARS-CoV or SARS-CoV-2 to bind to angiotensin converting enzyme 2 (ACE2), and S protein of MERS-CoV to dipeptidyl peptidase 4 (DPP4) (*17*), respectively, through their receptor-binding domain (RBD) (*18*). In addition to ACE2 and DPP4, some recent studies suggest that there are possible other coronavirus-associated receptors and factors that facilitate the infection of SARS-CoV-2 (*19*), including the cell surface proteins Basignin (BSG or CD147) (*20*), and CD209 (*21*). Recently, clinical data have revealed that SARS-CoV-2 can infect several organs where ACE2 expression could not be detected in healthy individuals (*22, 23*), which highlights the need of closer inspection of virus entry mechanisms.

The binding of S protein to a cell-surface receptor is not sufficient for infection of host cell (*24*). In the cell-surface membrane fusion mechanism, after binding to the receptor, the S protein requires proteolytic activation by cell surface proteases like TMPRSS2, TPMRSS4, or other members of the TMPRSS family (*14, 25, 26*), followed by the fusion of virus and target cell membranes. In the alternative receptor mediated endocytosis mechanism, the endocytosed virion is subjected to an activation step in the endosome, resulting in the fusion of virus and endosome membranes and the release of the viral genome into the cytoplasm. The endosomal cysteine proteases cathepsin B (CTSB) and cathepsin L (CTSL) (*27*) might be involved in the fusion of virus and endosome membranes. Availability of these proteases in target cells largely determines whether viruses infect the cells through cell-surface membrane fusion or receptor mediated endocytosis. How the presence of these proteases impacts efficiency of infection with SARS-CoV-2, SARS-CoV and MERS-CoV, still remains elusive.

When the virus enters a cell, it may trigger an innate immune response, a crucial component of the defense against viral invasion. Compounds that regulate innate immune responses can be introduced as antiviral agents (*10*). The innate immune system is initialized as pattern recognition receptors (PRRs) such as Toll-like receptors (TLRs) and cytoplasmic retinoic acid-inducible gene I (RIG-I) like receptors (RLRs) recognize molecular structures of the invading virus (*28, 29*). This pattern recognition activates several signaling pathways and then downstream transcription factors such as interferon regulator factors (IRFs) and nuclear factor *κ*B (NF-*κ*B). Transcriptional activation of IRFs and NF-*κ*B stimulates the expression of type I (*α* or *β*) and type III (*λ*) interferons (IFNs). IFN-*α* (IFNA1, IFNA2, etc), IFN-*β* (IFNB1) and IFN-*λ* (IFNL1-4) are important cytokines of the innate immune responses. IFNs bind and induce signaling through their corresponding receptors (IFNAR for IFN-*α*/*β* and IFNLR for IFN-*λ*), and subsequently induce expression of IFN-simulated genes (ISGs) (e.g. MX1, ISG15 and OASL) and pro-inflammatory chemokines (e.g. CXCL8 and CCL2) to suppress viral replication and dissemination (*30, 31*). Dysregulated inflammatory host response results in acute respiratory distress syndrome (ARDS), a leading cause of COVID-19 mortality (*32*).

One attractive therapy option to combat COVID-19 is to harness the IFN-mediated innate immune responses. Clinical trials with type I and type III IFNs for treatment of COVID-19 have been conducted and many more are still ongoing (*33, 34*). In this regard, the kinetics of the secretion of IFNs in the course of SARS-CoV-2 infection needs to be defined. Unfortunately, some results on the host innate immune responses to SARS-CoV-2 are apparently at odds with each other (*35–39*), e.g. it is unclear whether SARS-CoV-2 infection induces low IFNs and moderate ISGs (*35*), or robust IFN responses and markedly elevated expression of ISGs (*36–39*). This has to be clarified. The use of IFNs as a treatment in COVID-19 is now a subject of debate as well (*40*). Thus, the kinetics of IFN secretion relative to the kinetics of virus replication need to be thoroughly examined to better understand the biology of IFNs in the course of SARS-CoV-2 infection and thus provide guidance to identify the temporal window of therapeutic opportunity.

We have collected and analyzed a diverse set of publicly available transcriptome data (*35, 41–45*): (1) bulk RNA-Seq data with different types of cells, including human non-small cell lung carcinoma cell line (H1299), human lung fibroblast-derived cells (MRC5), human alveolar basal epithelial carcinoma cell line (A549), A549 cells transduced with a vector expressing human ACE2 (A549-ACE2), primary normal human bronchial epithelial cells (NHBE), heterogeneous human epithelial colorectal adenocarcinoma cells (Caco2), and African green monkey (*Chlorocebus sabaeus*) kidney epithelial cells (Vero E6) infected with SARS-CoV-2, SARS-CoV and MERS-CoV (Table 1); (2) RNA-Seq data of lung samples, peripheral blood mononuclear cell (PBMC) samples, and bronchoalveolar lavage fluid (BALF) samples of COVID-19 patients and their corresponding healthy controls (Table 1 and Table 2). Using this collection, we systemically evaluated the replication and transcription status of virus in these cells, expression levels of coronavirus-associated receptors and factors, as well as the innate immune responses of these cells during virus infection.

**Table 1.** Data of cell lines (cells) included in this study

| Virus | Virus strain | Virus dose (MOI) | Time | Replicates | Species of origin | Cell type | Library preparation | Accession number |
| --- | --- | --- | --- | --- | --- | --- | --- | --- |
| SARS-CoV-2 | USA-WA1/2020 | 2 | 24h | 3 | Homo sapiens | NHBE | polyA+ selection | GSE147507 |
| Mock | Mock | Mock | 24h | 3 | Homo sapiens | NHBE | polyA+ selection | GSE147507 |
| SARS-CoV-2 | USA-WA1/2020 | 0.2 | 24h | 3 | Homo sapiens | A549 | polyA+ selection | GSE147507 |
| Mock | Mock | Mock | 24h | 3 | Homo sapiens | A549 | polyA+ selection | GSE147507 |
| SARS-CoV-2 | USA-WA1/2020 | 2 | 24h | 3 | Homo sapiens | A549 | polyA+ selection | GSE147507 |
| Mock | Mock | Mock | 24h | 3 | Homo sapiens | A549 | polyA+ selection | GSE147507 |
| SARS-CoV-2 | USA-WA1/2020 | 0.2 | 24h | 3 | Homo sapiens | A549-ACE2 | polyA+ selection | GSE147507 |
| Mock | Mock | Mock | 24h | 3 | Homo sapiens | A549-ACE2 | polyA+ selection | GSE147507 |
| SARS-CoV-2 | USA-WA1/2020 | 2 | 24h | 3 | Homo sapiens | A549-ACE2 | polyA+ selection | GSE147507 |
| Mock | Mock | Mock | 24h | 3 | Homo sapiens | A549-ACE2 | polyA+ selection | GSE147507 |
| SARS-CoV-2 | USA-WA1/2020 | 2 | 24h | 3 | Homo sapiens | Calu3 | polyA+ selection | GSE147507 |
| Mock | Mock | Mock | 24h | 3 | Homo sapiens | Calu3 | polyA+ selection | GSE147507 |
| SARS-CoV-2 | Munich/BavPat1/2020 | 0.3 | 24h | 2 | Homo sapiens | Calu3 | rRNA-depletion | GSE148729 |
| Mock | Mock | Mock | 24h | 2 | Homo sapiens | Calu3 | rRNA-depletion | GSE148729 |
| SARS-CoV-2 | Munich/BavPat1/2020 | 0.3 | 24h | 2 | Homo sapiens | Calu3 | polyA+ selection | GSE148729 |
| Mock | Mock | Mock | 24h | 2 | Homo sapiens | Calu3 | polyA+ selection | GSE148729 |
| SARS-CoV-2 | Munich/BavPat1/2020 | 0.3 | 24h | 2 | Homo sapiens | Caco2 | polyA+ selection | GSE148729 |
| Mock | Mock | Mock | 24h | 2 | Homo sapiens | Caco2 | polyA+ selection | GSE148729 |
| SARS-CoV-2 | Munich/BavPat1/2020 | 0.3 | 24h | 2 | Homo sapiens | H1299 | polyA+ selection | GSE148729 |
| Mock | Mock | Mock | 36h^ | 2 | Homo sapiens | H1299 | polyA+ selection | GSE148729 |
| SARS-CoV-2 | USA-WA1/2020 | 0.3 | 24h | 2* | Chlorocebus sabaeus | Vero E6 | rRNA-depletion | GSE153940 |
| Mock | Mock | Mock | 24h | 3 | Chlorocebus sabaeus | Vero E6 | rRNA-depletion | GSE153940 |
| SARS-CoV | Frankfurt strain | 0.3 | 24h | 2 | Homo sapiens | Calu3 | polyA+ selection | GSE148729 |
| SARS-CoV | Frankfurt strain | 0.3 | 24h | 2 | Homo sapiens | Calu3 | rRNA-depletion | GSE148729 |
| SARS-CoV | Frankfurt strain | 0.3 | 24h | 2 | Homo sapiens | Caco2 | polyA+ selection | GSE148729 |
| SARS-CoV | Frankfurt strain | 0.3 | 24h | 2 | Homo sapiens | H1299 | polyA+ selection | GSE148729 |
| SARS-CoV | Urbani strain | 0.1 | 24h | 3 | Homo sapiens | MRC5 | polyA+ selection | GSE56189 |
| SARS-CoV | Urbani strain | 3 | 24h | 3 | Homo sapiens | MRC5 | polyA+ selection | GSE56189 |
| SARS-CoV | Urbani strain | 0.1 | 24h | 3 | Chlorocebus sabaeus | Vero E6 | polyA+ selection | GSE56189 |
| SARS-CoV | Urbani strain | 3 | 24h | 3 | Chlorocebus sabaeus | Vero E6 | polyA+ selection | GSE56189 |
| MERS-CoV | EMC/2012 | 0.1 | 24h | 3 | Homo sapiens | MRC5 | polyA+ selection | GSE56189 |
| MERS-CoV | EMC/2012 | 3 | 24h | 3 | Homo sapiens | MRC5 | polyA+ selection | GSE56189 |
| MERS-CoV | EMC/2012 | 0.1 | 24h | 3 | Chlorocebus sabaeus | Vero E6 | polyA+ selection | GSE56189 |
| MERS-CoV | EMC/2012 | 3 | 24h | 3 | Chlorocebus sabaeus | Vero E6 | polyA+ selection | GSE56189 |
| Mock | Mock | Mock | 24h | 3 | Homo sapiens | MRC5 | polyA+ selection | GSE56189 |
| Mock | Mock | Mock | 24h | 3 | Homo sapiens | Vero E6 | polyA+ selection | GSE56189 |

^No corresponding 24h mock control samples for H1299 cells, 36h mock control samples were used instead.
\* There are three replicates, but when the manuscript was in preparation only two of them are available for downloading.

**Table 2.** Data of COVID-19 patients included in this study

| Individuals | Tissue | Data Type | Accession number |
| --- | --- | --- | --- |
| 2 | bronchoalveolar lavage fluid from COVID-19 patients | bulk RNA-Seq | CRA002390 |
| 3 | bronchoalveolar lavage fluid from healthy negative control | bulk RNA-Seq | PRJNA434133^ |
| 3 | peripheral blood mononuclear cells from COVID-19 patients | bulk RNA-Seq | CRA002390 |
| 3 | peripheral blood mononuclear cells from healthy negative control | bulk RNA-Seq | CRA002390 |
| 2 | lung biopsy from postmortem COVID-19 patients | bulk RNA-Seq | GSE147507 |
| 2 | lung biopsy from healthy negative control | bulk RNA-Seq | GSE147507 |
| 6 | bronchoalveolar lavage fluid from COVID-19 patients (severe) | single cell RNA-Seq | GSE145926 |
| 3 | bronchoalveolar lavage fluid from COVID-19 patients (moderate) | single cell RNA-Seq | GSE145926 |
| 1 | bronchoalveolar lavage fluid from healthy negative control | single cell RNA-Seq | PRJNA526088* |

^Three samples under project PRJNA434133: SRR10571724, SRR10571730, and SRR10571732 were used.
\* One sample with accession number GSM3660650 under project PRJNA526088 was used.

## Results

### Different infection efficiency of SARS-CoV-2, SARS-CoV and MERS-CoV in different cell types

The RNA-Seq data for all samples can be aligned to the genome of the corresponding virus to evaluate the infection efficiency in cells, estimated by the mapping rate to the virus genome,i.e. the percentages of viral RNAs in intracellular RNAs. To assess the infection efficiency of SARS-CoV-2, SARS-CoV, and MERS-CoV in different types of cells, we collected and analyzed a comprehensive public datasets of RNA-Seq data of cells infected with these viruses at 24 hours post infection (hpi) with comparable multiplicity of cellular infection (MOI) (Table 1). MOI refers to the number of viruses that are added per cell in infection experiments. For example, if 2000 viruses are added to 1000 cells, the MOI is 2.

Our analysis shows that the infection efficiency of viruses can be both cell type dependent and virus dose dependent (Fig. 1). MERS-CoV can efficiently infect MRC5 and Vero E6 cells. However, the infection efficiency is influenced strongly by MOI in the same type of cells. Cells infected with low MOI, say 0.1, have significantly lower mapping rates than those with high MOI, say 3 (Fig. 1). For SARS-CoV and SARS-CoV-2, the infection efficiency is influenced strongly by cell type. For SARS-CoV-2, there is efficient virus infection in A549-ACE2, Calu3, Caco2, and Vero E6 cells, but not in A549, H1299, or NHBE cells (Fig. 1 and Table S1). The mapping rates in A549, H1299, and NHBE cells are low even at high MOIs (Fig. 1 and Table S1). Similar to SARS-CoV-2, the infection by SARS-CoV is also cell type dependent, Vero E6 cells and Calu3 cells show high mapping rates to SARS-CoV genome, but the mapping rates of SARS-CoV in MRC5 and H1299 cells are close to zero even at the high MOI of 3 (Fig. 1 and Table S1). Since “total RNA” (see Methods/Data collection) includes additional negative-strand templates of virus, the mapping rates are usually much higher than those that used the PolyA+ selection method in the same condition (Fig. 1 and Table S1).

**Fig. 1.**
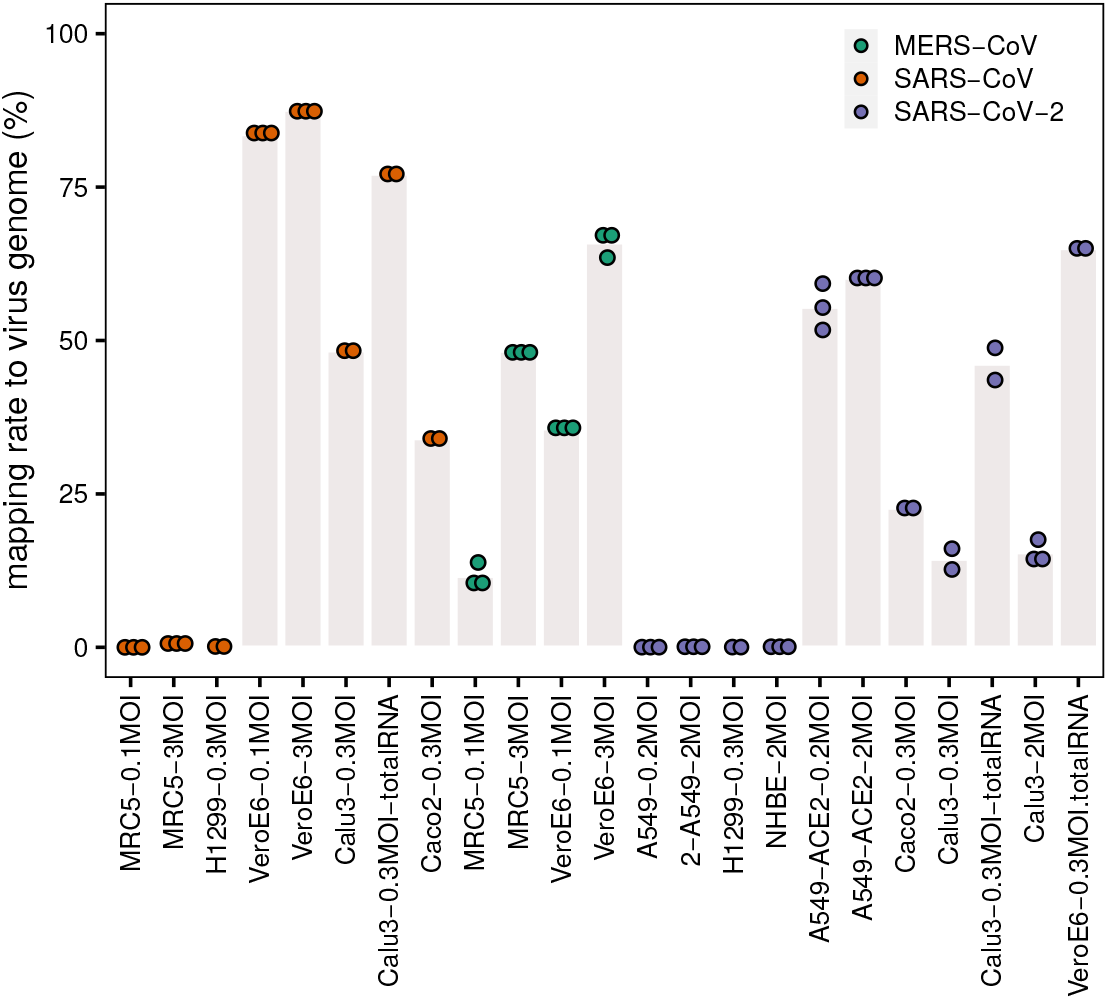
Mapping rate to virus genome. The dots represent the mapping rates to the virus genome for each individual replicate under the given conditions (cell line, MOI, and virus). Bar heights are mean mapping rates to the virus genome for each condition.

### Evidence for multiple entry mechanisms for SARS-CoV-2 and SARS-CoV

To examine the detailed replication and transcription status of these viruses in the cells, we calculated the number of reads (depth) mapped to each site of the corresponding virus genome (Fig. 2). For better comparison, these read numbers were log_10_ transformed. The replication and transcription of MERS-CoV, SARS-CoV-2 and SARS-CoV share an uneven pattern of expression along the genome, typically with a minimum depth in the first half of the viral genome, and the maximum towards the end. Among the parts with very high levels, there are especially coding regions for structural proteins, including S, E, M, and N proteins, as well as the first coding regions with nsp1 and nsp2. Interestingly, there is an exception for BALF samples in COVID-19 patients, which show a more irregular, fluctuating behavior along the genome (Fig. 2B). The deviation from the cellular expression pattern is not surprising because BALF is not a well-organized tissue but a mixture of many components, some of which will probably digest viral RNA.

**Fig. 2.**
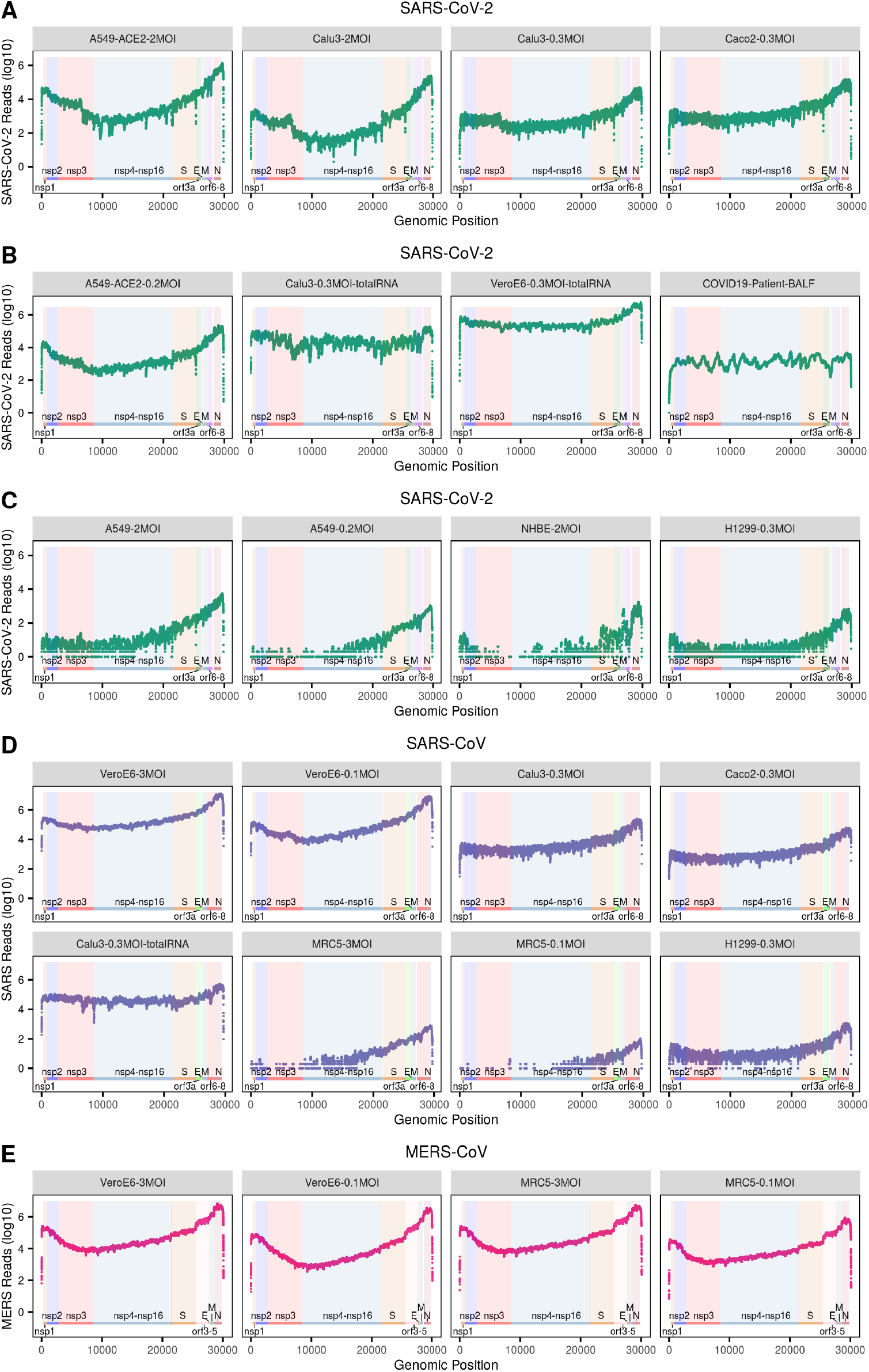
The number of reads mapped to the corresponding virus genome. (**A-E**) The dot plots show the number of reads mapped to each site of the corresponding virus genome. The annotation of the genome of each virus is from NCBI (SARS: GCF_000864885.1, SARS-CoV-2: GCF_009858895.2, MERS: GCF_000901155.1). Labels in grey title bars correspond to conditions as in Fig. 1.

Interestingly, the mentioned uneven transcription pattern of efficient infections with SARS-CoV-2, SARS-CoV, and MERS-CoV, is also visible for inefficient infection with SARS-CoV-2 in A549, NHBE, and H1299 cells, and SARS-CoV in H1299 and MRC5 cells (Fig. 2C, D), although there the total mapping rates to their corresponding virus genomes are much lower (Fig. 1).

To further elucidate the corresponding entry mechanisms for different types of cells, we examined the expression levels of those receptors and proteases that have already been described as facilitating target cell infection (Fig. 3).

**Fig. 3.**
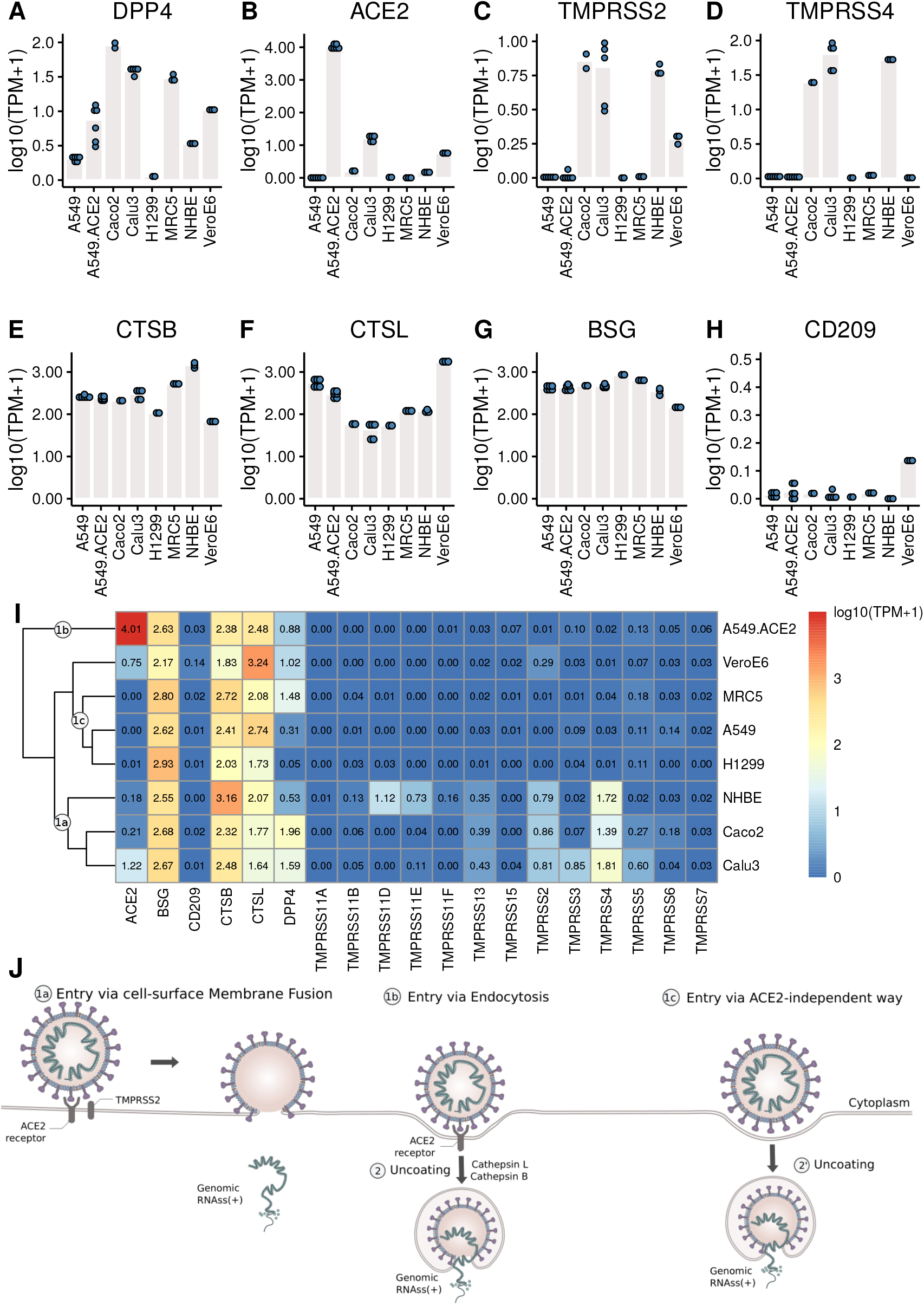
The expression levels of the receptors and proteases. (**A-H**) Each dot represents the expression value in each sample. (**I**) Heatmap of the expression levels of coronavirus associated **receptors** and factors of different cell types. Labels 1a, 1b, 1c mark cell clusters that likely share entry routes sketched in panel **J**. (**J**) Entry mechanisms involved in SARS-CoV-2 entry into cells. Schematic is based on a figure by Vega Asensio - Own work, CC BY-SA 4.0, https://commons.wikimedia.org/w/index.php?curid=88682468.

Our analysis shows that MERS-CoV can efficiently infect MRC5 and Vero E6 cells (Fig. 1 and Fig. 2E) that both express DPP4 (Fig. 3A), though compared to Vero E6 cells, MRC5 cells infected with MERS-CoV have higher expression levels of DPP4 (Fig. 3A), but lower mapping rates to the virus genome (Fig. 1). These observations show that higher expression levels of the receptor (DPP4) do not guarantee higher MERS-CoV infection efficiency in cells. This is also true for SARS-CoV-2 receptor ACE2, which is expressed three orders of magnitudes higher in A549-ACE2 cells than in Vero E6 cells (Fig. 3B), while both cells produce about the same amount of virus (Fig. 1).

Although SARS-CoV-2 can efficiently infect A549-ACE2 cells (Fig. 1 and Fig. 2), there is no expression of TMPRSS2 or TMPRSS4 (Fig. 3C, D), needed for the canonical cell-surface membrane fusion mechanism (Fig. 3J). However, there are considerable expression levels of CTSB and CTSL (Fig. 3E, F), which are involved in endocytosis (Fig. 3J).

In A549, H1299, and MRC5 cells, which do express small amounts of SARS-CoV-2 and SARS-CoV virus (Fig. 1, Fig. 2C, D), there is no ACE2 expression at all (Fig. 3B). This could point to an alternative ACE2-independent entry mechanism for SARS-CoV-2 and SARS-CoV (Fig. 3J). Since there were already reports about alternative SARS-CoV-2 receptors such as BSG/CD147 and CD209 (*20,21*), we examined their expressions in these cells as well (Fig. 3G, H). For all cells, the expression of BSG is at the same level of 2-3 (Fig. 3G), and the expression of CD209 is very low. Certainly, CD209 and BSG alone cannot explain the differences in virus expression (Fig. 1), nor can we exclude other low efficiency entry mechanisms. It could e.g. be that relatively inefficient alternative entry paths are often present but in some cells masked by more efficient entry via ACE2/TPMRSS.

To gain a comprehensive overview we clustered cells with respect to gene expression levels of coronavirus-associated receptors and factors (Fig. 3I), and summarized conceivable mechanisms accordingly (Fig. 3J). Since all cells show high expression levels of CTSB and CTSL, the major differences between these cells lie in the expression levels of ACE2, TMPRSS2 and TPMRSS4.

Cell-surface membrane fusion (Fig. 3J, 1a) might be mainly used in SARS-CoV-2 infection of Calu3, Caco2, and NHBE cells where there are low to moderate expression of ACE2 and moderate expression of TMPRSS2 and TMPRSS4. Endocytosis (Fig. 3J, 1b) might be mainly used in SARS-CoV-2 infection of A549-ACE2 cells where ACE2 is expressed at high levels but there is no expression of TMPRSS2 or TMPRSS4. An alternative ACE2-independent way (Fig. 3J, 1c) in absence of ACE2, TMPRSS2, or TMPRSS4 could be mainly employed in SARS-CoV-2 infection of MRC5, A549, and H1299 cells. Note that although the expression pattern of coronavirus-associated receptors and factors of NHBE cells is similar to that in Caco2 cells, NHBE cells are not infected efficiently by SARS-CoV-2. Vero E6 cells have moderate expression of ACE2, and low expression of TMPRSS2 and TMPRSS4, so all these entry mechanisms mentioned above could contribute to SARS-CoV-2 infection of Vero E6 cells.

### Strength of IFN/ISG response varies between cell lines and viruses, with strong response to SARS-CoV-2 in relevant cells

As a virus enters a cell, it may trigger an innate immune response, i.e. the cell may start expression of various types of innate immunity molecules at different strengths. There is currently an intense debate about which of these molecules, especially IFNs and ISGs, are expressed how strongly (*35–39*). We therefore focused in our analysis on innate immunity molecules such as IFNs, ISGs, and pro-inflammatory cytokines. To broaden the basis for conclusions, we analyzed, apart from cell lines, bulk RNA-Seq data of lung, PBMC, and BALF samples of COVID-19 patients, and single-cell RNA-Seq data of BALF samples from moderate and severe COVID-19 patients; for each type of patient data, we also included healthy controls. Gene expressions were compared quantitatively in terms of TPM (transcripts per million), as well as log fold changes (logFC) with respect to healthy controls (human samples) or mock-infected cultures (cell lines) (Fig. S1, Fig. S2).

The heatmap and clustering dendrogram of the logFC of IFNs, ISGs and pro-inflammatory cytokines in Fig. 4A reveal broadly two groups of samples with fundamentally different expression of ISGs, IFNs, and pro-inflammatory cytokines.

**Fig. 4.**
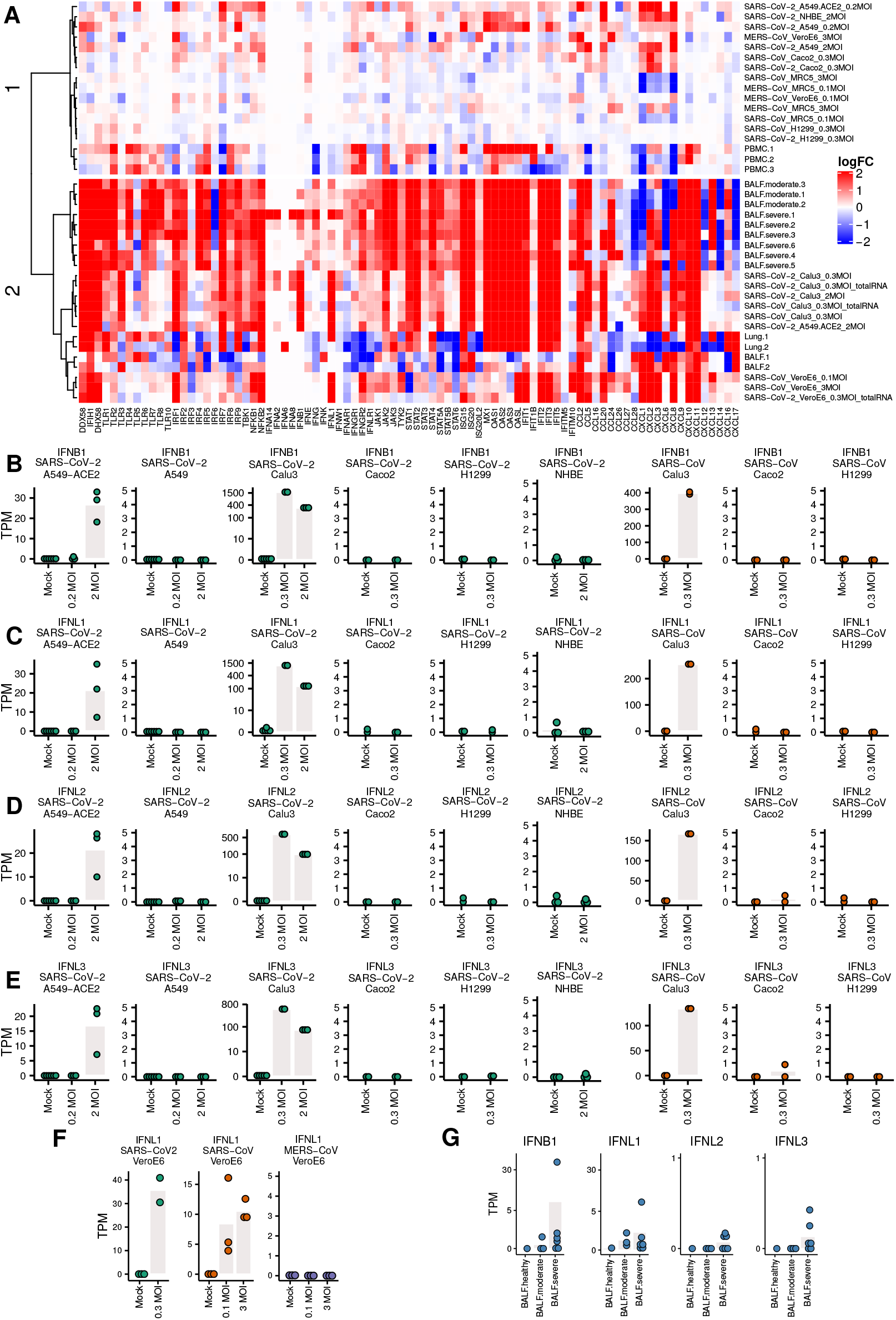
Expression levels of genes related to immune responses. (**A**) Heatmap of the logFC of IFNs, ISGs and pro-inflammatory cytokines. The clustering of samples produces a cluster 1 (top) with little IFN/ISG expression comprising MERS infections and non-infectable cells/SARS-CoV-1/2 (except for Caco2 cells), and a cluster 2 (bottom) strong IFN/ISG expression with SARS-CoV-1/2 infectable cells and patient samples. (**B-G**) Expression levels of IFNs. Each dot represents the expression value of a sample. Bars indicate mean expression levels (in TPM) of respective IFN at different MOI values.

The top cluster in Fig. 4A are samples that show weaker innate immune response, including the two PBMC samples of COVID-19 patients, A549, NHBE, Caco2, and H1299 cells infected with SARS-CoV-2 and A549-ACE2 cells infected with SARS-CoV-2 at lower MOI (0.2), MRC5 cells infected with SARS, MRC5 and Vero E6 cells infected with MERS. The bottom cluster in Fig. 4A are samples that show stronger innate immune response, including BALF and lung samples of COVID-19 patients, Calu3 cells infected with SARS-CoV-2, A549-ACE2 cells infected with SARS-CoV-2 at higher MOI (2), as well as Vero E6 cells infected with SARS-CoV-2 and SARS. Most of the samples in the bottom part show markedly elevated levels of ISGs and elevated pro-inflammatory cytokines. An exception in the bottom cluster are four samples, namely Lung.1/2 and BALF.1/2, with a mixture of up- and down-regulation of ISGs and pro-inflammatory cytokines. In this respect, these four samples from patients with unknown COVID-19 severity differ from the BALF samples from moderate and severe COVID-19 patients.

The expression levels of IFNs are not upregulated either in most of these lung, PBMC and BALF samples of COVID-19 patients where no information about the severity of infection of these COVID-19 patients are available. However, we estimated the severity of their infection by aligning all the samples to SARS-COV-2 virus genome. There are no (0.00%) reads mapping to the SARS-CoV-2 genome in the PBMC samples. For the two BALF samples, there are low mapping rates (1.56% and 0.65%) to SARS-CoV-2 genome. The expression levels of ACE2 in these tissues (PBMC, lung and BALF samples) of healthy individuals are around zero (Fig. S8), which explains why there are almost no virus reads in these tissues.

One of the two lung samples (accession number: SAMN14563387) has slightly upregulated IFNL1 (Fig. S6), which had been ignored in the original publication (*35*), although the total mapping rates to virus genome are both 0.00% for these two lung samples. We then checked the detailed coverage along the virus genome. There were a small number of virus reads aligned to SARS-CoV-2 genome in this sample (Fig. S7). Different from other lung samples that did not express ACE2, this lung sample expressed ACE2 at a considerable level (5.45 TPM, Table S2). This result implies that when SARS-CoV-2 enters into lung successfully, or when the lung tissue chosen for sequencing are successfully infected by SARS-CoV-2, IFNs (at least IFNL1) can be upregulated.

Calu3 cells infected with SARS-CoV and SARS-CoV-2, and A549-ACE2 cells infected with SARS-CoV-2 at a high MOI of 2 have upregulated IFNB1, IFNL1, IFNL2 and IFNL3 (Fig. 4B-E). A549, H1299, NHBE (Fig. 4B-E), and MRC5 cells (Fig. S3), which do not support efficient virus infection, show no upregulation of IFNs. Low levels of IFN expression are also observed in Caco2 cells, which are efficiently infected with SARS-CoV and SARS-CoV-2. The same is true for A549-ACE2 cells infected with SARS-CoV-2 at low MOI of 0.2. In Vero E6 cells IFNL1 is upregulated as well in infected with SARS-CoV and SARS-CoV-2, but not with MERS-CoV (Fig. 4F). In BALF samples of moderate and severe COVID-19 patients, upregulation of IFNs was not as obvious as in Calu3 cells, but is still present in some patients. These observations demonstrate that the innate immune response depends in complex ways on cell line, viral dose, and virus.

Several studies (*36–39*) reported robust IFN responses and markedly elevated expression of ISGs in SARS-CoV-2 infection of different cells and patient samples. Conversely, the study by (*35*) concluded that weak IFN response and moderate ISG expression are characteristic for SARS-CoV-2 infection. This apparent contradiction can be resolved if we consider that Ref. (*35*) generalized from patient samples and cells that were only weakly infected, and that in such cases the host, in fact, responds with low levels of IFNs and ISGs. On the other hand, Ref. (*35*) treated efficiently infected cells, such as Calu3 and A549-ACE2 (at MOI 0f 2) as exceptions. However, our meta-analysis shows that these are not exceptions but typical for severely infected target cells that have robust IFN responses and ISG expressions (cluster 2 in Fig. 4A).

## Discussion

One attractive potential anti-SARS-CoV-2 therapy is intervention in the cell entry mechanisms (*12*). However, the entry mechanisms of SARS-CoV-2 into human cells are partly unknown. During the last few months scientists have confirmed that SARS-CoV-2 and SARS-CoV both use human ACE2 as entry receptor, and human proteases like TMPRSS2 and TMPRSS4 (*8, 14, 25*), and lysosomal proteases like CTSB and CTSL (*27*) as entry activators. Since ACE2 is beneficial in cardiovascular diseases such as hypertension or heart failure (*46*), treatments targeting ACE2 could have a negative effect. Inhibitors of CTSL (*47*) or TMPRSS2 (*14*) are seen as potential treatment options for SARS-CoV and SARS-CoV-2. However, recently alternate coronavirus-associated receptors and factors including BSG/CD147 (*20*) and CD209 (*21*) have been proposed to facilitate virus invasion. Additionally, clinical data of SARS-CoV-2 infection have shown that SARS-CoV-2 can infect several organs where ACE2 expression could not be detected (*22, 23*), urging us to explore other potential entry routes.

First, our analyses here have shown that even without expression of TMPRRS2 or TM-PRSS4, high SARS-CoV-2 infection efficiency in cells is possible (Fig. 1A, C) with considerable expression levels of CTSB and CTSL (Fig. 2E, F). This suggests receptor mediated endocytosis (*15, 16, 27*) as an alternative major entry mechanism. Given this TMPRSS-independent route, TMPRSS inhibitors will likely not provide complete protection. The studies designed to predict the tropism of SARS-CoV-2 by profiling the expression levels of ACE2 and TMPRSS2 across healthy tissues (*48, 49*) may need to be reconsidered as well.

Second, the evidence presented in our study suggests further, possibly undiscovered entry mechanism for SARS-CoV-2 and SARS-CoV (Fig. 2). Although BSG/CD147 has been recently proposed as an alternate receptor (*20*), later experiments reported there was no evidence supporting the role of BSG/CD147 as a putative spike-binding receptor (*50*). The expression patterns of BSG/CD147 in different types of cells observed in our study could not explain the difference in virus loads observed in these cells either. CD209 and CD209L were recently reported as attachment factors to contribute to SARS-CoV-2 infection in human cells as well (*21*). However, CD209 expression in the cell lines included here is low. Another reasonable hypothesis could be that the inefficient ACE2-independent entry mechanism we observed could be macropinocytosis, one endocytic pathway that does not require receptors (*51*). Until now there is still no direct evidence for macropinocytosis involvement in SARS-CoV-2 and SARS-CoV entry mechanism. To confirm such an involvment, specific experiments are needed. Moreover, this ACE2-independent entry mechanism, only enables inefficient infection by SARS-CoV and SARS-CoV-2 (Fig. 2) and therefore cannot be a major entry mechanism.

Fig. 3J summarizes the outcomes of our study with respect to entry mechanisms. The observations with the broad range of transcriptome data can only be explained if there are several entry routes. This is certainly a challenge to be reckoned with in the development of antiviral therapeutics (*52*).

Another attractive potential anti-SARS-CoV-2 point of attack is supporting the human innate immune system to kill the infected cells and, thus disrupt viral replication. Not surprisingly, research in this area is flourishing but sometimes generates conflicting results, especially on the involvement of type I and III IFNs and ISGs (*35–39*). The results of our analyses could help to dissolve the confusion on the involvement of IFNs and ISGs.

We found that immune responses in Calu3 cells infected with SARS-CoV and SARS-CoV-2 resemble those of BALF samples of moderate and severe COVID-19 patients, with elevated levels of type I and III IFNs, robust ISG induction as well as markedly elevated pro-inflammatory cytokines, in agreement with recent studies (*36–39*). However this picture differs from the one reported by (*35*) with low levels of IFNs and moderate ISGs. This latter study was partially based on A549 cells and NHBE cells with nearly no ACE2 expression and very low mapping rate to the viral genome, and lung samples of two patients (both show 0.00% mapping rate to virus genome). Hence, given that there was no efficient virus infection in theses cells, the low levels of IFNs and ISGs were to be expected. However, in one of the lung samples sequenced by (*35*) (accession number: SAMN14563387), we observed a slight upregulation of IFNL1 (Fig. S6), which was ignored in the original publication, together with considerable ACE2 expression (Table S2) (5.45 TPM), and a few virus reads aligned to SARS-CoV-2 genome (Fig. S7). This results suggests that levels of IFNs are ISGs are associated with viral load and severity of virus infection.

We found low induction of IFNs and moderate expression of ISGs in PBMC samples and BALF samples of COVID-19 patients (Fig. 4, Fig. S5). In these PBMC samples, there are no (0.00%) virus reads mapping to the SARS-CoV-2 genome. The failure to detect virus reads in these three PBMC samples can be explained by the absence of efficient entry routes (e.g. no expression of ACE2 in PBMC samples of healthy individuals, Fig. S8), or with the cell types being otherwise incompatible with viral replication. This observation is consistent with the studies on SARS-CoV (*53–55*) with abortive infections of macrophages, monocytes, and dendritic cells; moreover, replication of SARS-CoV in PBMC samples is also self-limiting. However, due to the limited number of PBMC, BALF and lung samples included in this study, and the lack of the information of infection stage and infection severity of these COVID-19 patients, the assessment of IFNs and ISGs as well as the infection of SARS-CoV-2 in these samples may not be representative of host response against SASR-CoV-2. Future studies that include also other affected organs of more patients with different infection stages and severity are necessary for a better understanding of the immune responses.

Several unexpected observations need further investigations. First, A549-ACE2 and Caco2 cells are efficiently infected with low MOI of 0.2 and 0.3, respectively, (Fig. 1), but fail to upregulate INF expression (Fig. 4B-E). Their cellular immune responses are more similar to those of cells that cannot support efficient virus infection (Fig. 4A). These results suggest that in Caco2 and A549-ACE2 cells the invasion of SARS-CoV-2 or SARS-CoV at low MOI shuts down or fails to activate the innate immune system.

Based on the results observed above, multiple factors including disease severity, different organs, cell types and virus dose contribute to the variability in the innate immune responses. For a better characterization of the innate immune responses, a more comprehensive profiling is necessary, including of patients with infections in different stages, different levels of severity, and different clinical outcomes of the infection. Further, a larger array of cell types should be profiled over time after infection with different virus doses. In this way we would be better able to understand the kinetics of IFNs and ISGs in response to SARS-CoV-2 infection.

In summary, our study has comparatively analyzed an extensive data collection from different cell types infected with SARS-CoV-2, SARS-CoV and MERS-CoV, and from COVID-19 patients. We have presented evidence for multiple SARS-CoV-2 entry mechanisms. We could also dissolve apparent conflicts on innate immune responses in SARS-CoV-2 infection (*35–39*), by drawing upon a larger set of cell types and infection severity. The results emphasize the complexity of interactions between host and SARS-CoV-2, offer new insights into pathogenesis of SARS-CoV-2, and can inform development of antiviral drugs.

## Materials and Methods

### Data collection

After the successful release of the virus genome into the cytoplasm, a negative-strand genomic-length RNA is synthesized as the template for replication. Negative-strand subgenome-length mRNAs are formed as well from the virus genome as discontinuous RNAs, and used as the templates for transcription. In the public data we collected for the analysis, there are two main library preparation methods to remove the highly abundant ribosomal RNAs (rRNA) from total RNA before sequencing. One is polyA+ selection, the other is rRNA-depletion (*56*). It is known that coronavirus genomic and subgenomic mRNAs carry a polyA tail at their 3’ ends, so in the polyA+ RNA-Seq, we have (1) virus genomic sequence from virus replication, i.e. replicated genomic RNAs from negative-strand as template, and (2) subgenomic mRNAs from virus transcription; in the rRNA-depletion RNA-Seq we have (1) virus genomic sequence from virus replication: both replicated genomic RNAs from negative-strand as template and the negative-strand templates themselves, and (2) subgenomic mRNAs from virus transcription. PolyA+ selection was used if not specifically stated in this study, “total RNA” is used to specify that the rRNA-depletion method was used to prepare the sequencing libraries.

The raw FASTQ data of different cell types infected with SARS-CoV-2, SARS-CoV and MERS-CoV, and lung samples of COVID-19 patients and healthy controls were retrieved from NCBI (*57*) (https://www.ncbi.nlm.nih.gov/) and ENA (*58*) (https://www.ebi.ac.uk/ena) (accession numbers GSE147507 (*35*), GSE56189, GSE148729 (*41*) and GSE153940 (*59*)). The raw FASTQ data of PBMC and BALF samples of COVID-19 patients and corresponding controls were downloaded from BIG Data Center (*60*) (https://bigd.big.ac.cn/) (accession number CRA002390) (*42*), and the raw FASTQ data for BALF healthy control samples were downloaded from NCBI (accession numbers SRR10571724, SRR10571730, and SRR10571732 under project PRJNA434133 (*43*)). The preprocessed single cell RNA-Seq data of BALF samples from 6 severe COVID-19 patients and 3 moderate COVID-19 patients were downloaded from NCBI with accession number GSE145926 (*44*). The preprocessed single cell RNA-Seq data of BALF sample from a healthy control was retrieved from NCBI (accession number GSM3660650 under project PRJNA526088 (*45*)). Detailed information about these public datasets are available in the supplementary file: Supplementary.pdf

For analysis, the human GRCh38 release 99 transcriptome and the green monkey (*Chlorocebus sabaeus*) ChlSab1.1 release 99 transcriptome and their corresponding annotation GTF files were downloaded from ENSEMBL (*61*) (https://www.ensembl.org). The reference virus genomes were downloaded from NCBI: SARS-CoV-2 (GenBank: MN985325.1), SARS-CoV (GenBank: AY278741.1), MERS-CoV (GenBank: JX869059.2).

### Data analysis workflow

The workflow of this study is summarized in Fig. S1 and Fig. S2 in the supplementary file: Supplementary.pdf. The quality of the raw FASTQ data was examined with FastQC (*62*). Trimmomatic-0.36 (*63*) was used to remove adapters and filter out low quality reads with parameters “-threads 4 -phred33 ILLUMINACLIP:adapters.fasta:2:30:10 HEADCROP:10 LEADING:20 TRAILING:20 SLIDINGWINDOW:4:20 MINLEN:36”. The clean RNA sequencing reads were then pseudo-aligned to reference transcriptome and quantified using Kallisto (version 0.43.1) (*64*) with parameters “-b 30 –single −l 180 -s 20” for single-end sequencing data and with parameter “-b 30” for paired-end sequencing data. Expression levels were calculated and summarized as transcripts per million (TPM) on gene levels with Sleuth (*65*), and logFC was then calculated for each condition. The single cell RNA-Seq data were summarized across all cells to obtain “pseudo-bulk” samples. R packages EDASeq (*66*) and org.Hs.eg.db (*67*) were used to obtain gene length, and TPM was calculated with the “calculateTPM” function of R package scater (*68*). logFC was then calculated for each patient.

The clean RNA-Seq data were also aligned to the virus genome with Bowtie 2 (*69*) (version 2.2.6) and the aligned BAM files were created, and the mapping rates to the virus genomes were obtained as well. SAMtools (*70*) (version 1.5) was then used for sorting and indexing the aligned BAM files. The “SAMtools depth” command was used to produce the number of aligned reads per site along the virus genome.

The heatmap in Fig. 3I was made by pheatmap R package (*71*), “complete” clustering method was used for clustering the rows and “euclidean” distance was used to measure the cluster distance. The heatmap in Fig. 4A was made by ComplexHeatmap R package (*72*). “complete” clustering method was used for clustering the rows and columns and “euclidean” distance was used to measure the cluster distance.

## Supporting information

Including supplemental methods and figures

## Acknowledgements

The authors thank professor Ke Xu from Wuhan University and professor Dimitri Lavillette from Institut Pasteur of Shanghai for helpful conversations.

## Funding

This work was partially funded by grant 01Kl20185B (SECOVIT) of the German Federal Ministry of Education and Research.

## Author Contributions

Pei Hao and Yingying Cao conceived the research. Daniel Hoffmann, Pei Hao, and Yingying Cao designed the analyses. Yingying Cao, Xintian Xu conducted the analyses. All authors wrote the manuscript.

## Competing Interests

The authors declare that they have no competing financial interests.

## Data and materials availability

Additional data and materials are available online.

## Notes

### Competing Interest Statement

The authors have declared no competing interest.

### Summary of Updates

author affiliation updated

