## Supplementary material for "Comprehensive comparison of transcriptomes in SARS-CoV-2 infection: alternative entry routes and innate immune responses": Including supplemental methods and figures

### 1 Additional information about public data

All data can be downloaded from public repositories, the three main sources are NCBI (*1*) (https://www.ncbi.nlm.nih.gov/) and ENA (*2*) (https://www.ebi.ac.uk/ena) and BIG Data Center (*3*) (https://bigd.big.ac.cn/).

#### 1.1 GSE147507 dataset (*4*)

From this dataset we downloaded: Biological triplicates of primary human lung epithelium (NHBE) which were mock treated or infected with SARS-CoV-2 (USA-WA1/2020) at an MOI of 2; Biological triplicates of transformed lung alveolar (A549) cells which were mock treated or infected with SARS-CoV-2 (USA-WA1/2020) at an MOI of 0.2 or 2; Biological triplicates of transformed lung alveolar (A549) transduced with a vector expressing human ACE2, which were also mock treated or infected with SARS-CoV-2 (USA-WA1/2020) at an MOI of 0.2 or 2; Biological triplicates of transformed lung-derived Calu-3 cells which were mock treated or infected with SARS-CoV-2 (USA-WA1/2020) at an MOI of 2; COVID-19 patient samples: Uninfected human lung biopsies derived from one male (age 72) and one female (age 60) and used as control biological replicates, and lung samples derived from a single male COVID-19 deceased patient (age 74) which were processed in technical replicates. Library preparation method polyA+ selection was used to remove rRNAs before sequencing.

#### 1.2 GSE148729 dataset (*5*)

From this dataset we downloaded biological replicates of Calu-3, Caco-2 and H1299 cells which were mock treated or infected with SARS-CoV-2 (patient isolate BetaCoV/Munich/BavPat1/2020/EPI_ISL_406862) or SARS-CoV (Frankfurt strain) at an MOI of 0.3. Library preparation method polyA+ selection was used to remove rRNAs before sequencing Caco-2 and H1299 cells. For Calu-3 cells, two library preparation method polyA+ selection and rRNA-depletion were used respectively to remove rRNAs before sequencing.

#### 1.3 GSE153940 dataset

From this dataset we downloaded RNA sequencing data of Vero E6 cells which were either mock-infected or infected with SARS-CoV-2 USA-WA1/2020 (MOI = 0.3) with three replicates. However, when we downloaded the data one sample with accession number GSM4658806 was not available for downloading. Cells were harvested at 24 hours after infection, and rRNA-depletion method was used to extract RNA for sequencing.

#### 1.4 GSE56189 dataset

From this dataset we downloaded: Biological triplicates of MRC5 and Vero E6 cells which were mock treated or infected with SARS-CoV (Urbani strain) or MERS-CoV (EMC/2012) at an MOI of 0.1 or 3. Library preparation method polyA+ selection was used to remove rRNAs before sequencing.

#### 1.5 CRA002390 dataset (*6*)

This dataset is public available in https://bigd.big.ac.cn/gsa/browse/CRA002390. From this dataset we downloaded: The raw FASTQ data of PBMC and BALF samples of COVID-19 patients and corresponding PBMC controls.

#### 1.6 PRJNA434133 dataset (*7*)

From this dataset we downloaded the raw FASTQ data for BALF healthy control samples with accession numbers SRR10571724, SRR10571730, and SRR10571732.

#### 1.7 GSE145926 dataset (*8*)

From this dataset we downloaded the preprocessed single cell RNA-Seq data of BALF samples from 6 severe COVID-19 patients and 3 mild COVID-19 patients.

#### 1.8 PRJNA526088 dataset (*9*)

From this dataset we downloaded the preprocessed single cell RNA-Seq data of BALF sample from a healthy control with accession number GSM3660650.

### 2 Supplementary figures

#### Fig. S1:

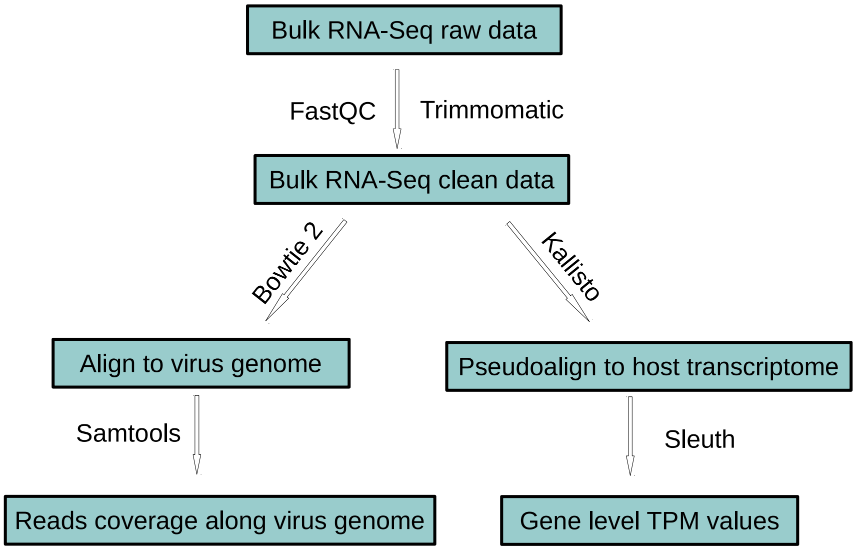


#### Fig. S2:

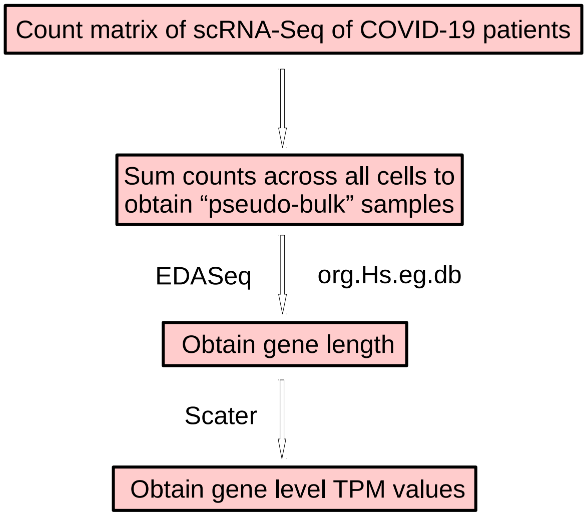

**Fig. S3:**

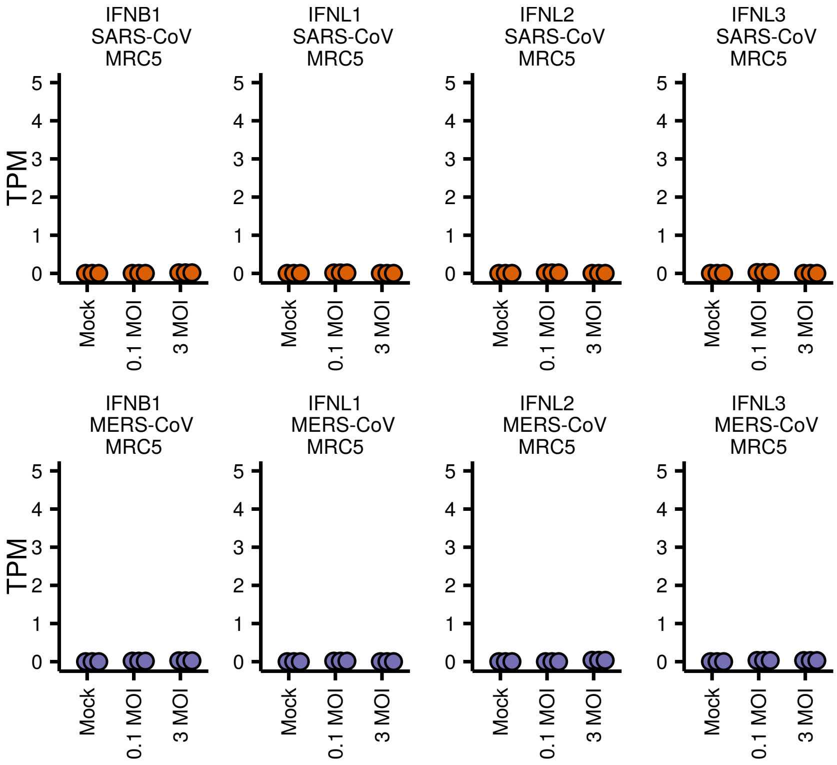
IFNB1
SARS-CoV
MRC5
IFNL1
SARS-CoV
MRC5
IFNL2
SARS-CoV
MRC5
IFNL3
SARS-CoV
MRC5
IFNB1
MERS-CoV
MRC5
IFNL1
MERS-CoV
MRC5
IFNL2
MERS-CoV
MRC5
IFNL3
MERS-CoV
MRC5
TPM
0
1
2
3
4
5
Mock
0.1 MOI
3 MOI

**Fig. S4:**

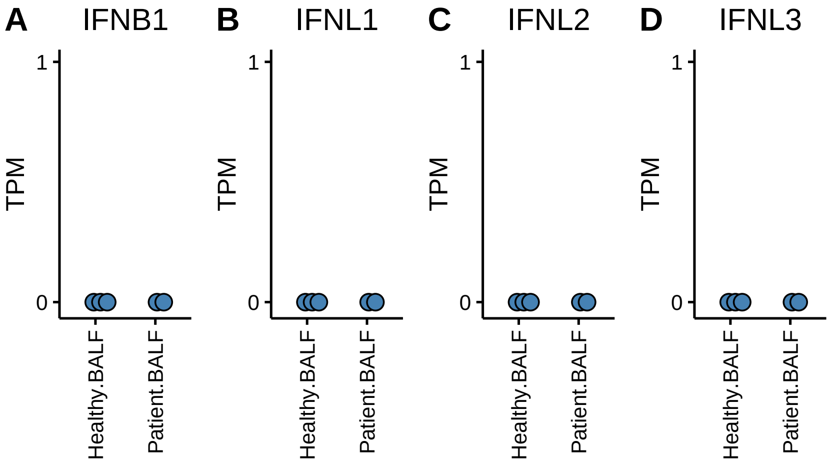

A
IFNB1
B
IFNL1
C
IFNL2
D
IFNL3
TPM
1
0
Healthy.BALF
Patient.BALF

**Fig. S5:**

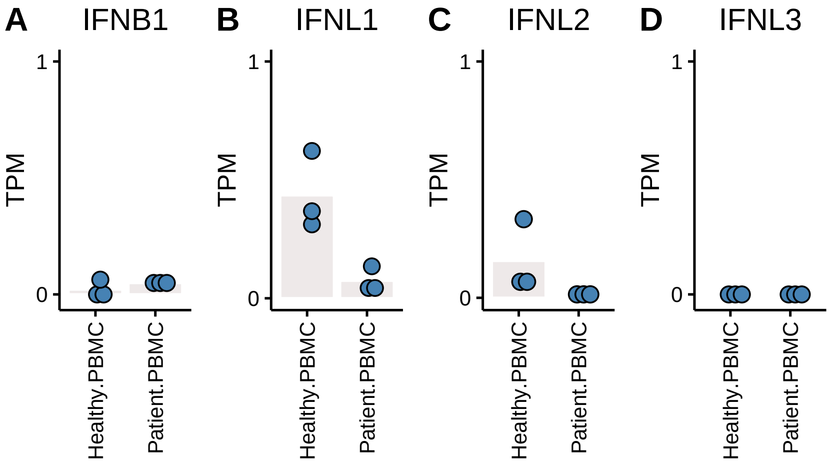
A
IFNB1
B
IFNL1
C
IFNL2
D
IFNL3
TPM
1
0
Healthy.PBMC
Patient.PBMC

**Fig. S6:**

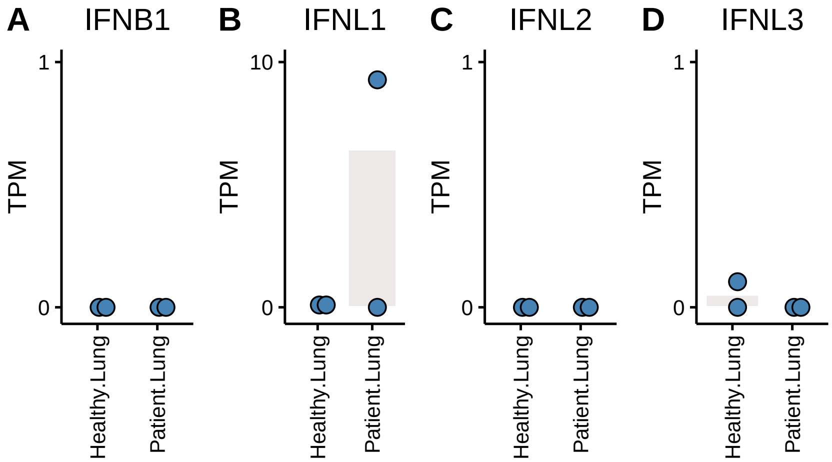
A
IFNB1
B
IFNL1
C
IFNL2
D
IFNL3
TPM
1
0
10
Healthy.Lung
Patient.Lung

**Fig. S7:**

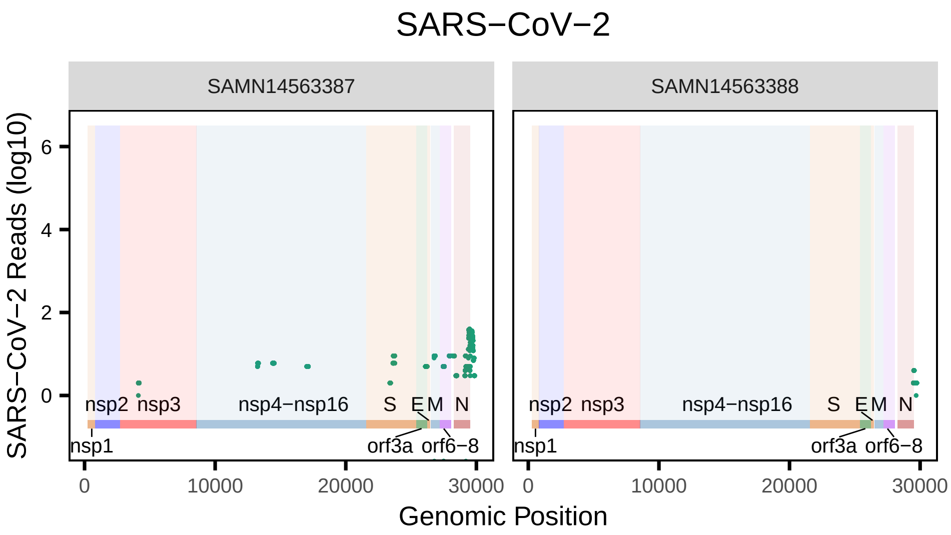
SARS−CoV−2
SAMN14563387
SAMN14563388
SARS−CoV−2 Reads (log10)
0
2
4
6
nsp1
nsp2
nsp3
nsp4−nsp16
S
E
M
N
orf3a
orf6−8
0
10000
20000
30000
Genomic Position

**Fig. S8:**

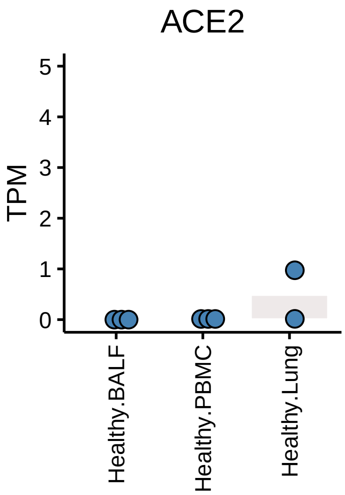
